# Natural IgA and *TNFRSF13B* polymorphism: a double edged sword fueling balancing selection

**DOI:** 10.1101/2021.01.29.428850

**Authors:** Jeffrey L. Platt, Mayara Garcia de Mattos Barbosa, Daniel Huynh, Adam R. Lefferts, Juhi Katta, Cyra Kharas, Peter Freddolino, Christine M. Bassis, Christiane Wobus, Raif Geha, Richard Bram, Gabriel Nunez, Nobuhiko Kamada, Marilia Cascalho

## Abstract

*TNFRSF13B* encodes the “transmembrane-activator and CAML-interactor” (TACI) receptor, which drives plasma cell differentiation. Although *TNFRSF13B* supports host defense, dominant-negative *TNFRSF13B* alleles are common in humans and other species and only rarely associate with disease. We reasoned the high frequency of disruptive *TNFRSF13B* alleles reflects balancing selection, the loss of function conferring advantage in some settings. Testing that concept, we asked whether and how a common human dominant negative variant, TNFRSF13B A181E, imparts resistance to enteric pathogens. Mice engineered to express mono-allelic or bi-allelic A144E variants of *tnrsf13B*, corresponding to A181E exhibited striking resistance to pathogenicity and transmission of *C. rodentium*, a murine pathogen that models enterohemorrhagic *E. coli*, and resistance was principally owed to deficiency of natural IgA in the intestine. In wild type mice with gut IgA and in mutant mice fed IgA, binding of Ig induces expression of *LEE* encoded virulence genes, which confer pathogenicity and transmission*. C. rodentium* and probably some other enteric organisms thus appropriated binding of otherwise protective antibodies to signal induction of the virulence program and the high prevalence of *TNFRSF13B* dominant negative variants thus reflects balancing selection.

## Introduction

*TNFRSF13B* encodes the “transmembrane activator and CAML interactor” (TACI), a member of the TNF receptor superfamily and has been considered vital to immune fitness. TACI is the receptor for the B cell activating factor (BAFF) and “a proliferation induced ligand” (APRIL). Binding of BAFF or APRIL to TACI activates BLIMP-1 (1), the transcription factor that governs differentiation of B lymphocytes into plasma cells (2–4). Common variable immunodeficiency and IgA deficiency in humans and mice has been associated with *TNFRSF13B* null and dominant negative mutations (5, 6).

Since *TNFRSF13B* in humans supports a key facet of immune fitness, it might be surprising that surveys of normal populations reveal extraordinary polymorphism - 951 *TNFRSF13B* missense and only 383 synonymous mutations (https://useast.ensembl.org/index.html) - and a high frequency of dominant negative alleles (7). Indeed, most individuals with TNFRSF13B variants that disrupt function are healthy (7, 8). Nor is this paradoxical diversity limited to humans, as *TNFRSF13B* polymorphism also occurs in other species. For example, 17 missense, 2 stop gained and 2 splice variants have been reported in mice (https://useast.ensembl.org/index.html) (9). The mechanism underlying *TNFRSF13B* polymorphism is unknown, although missense alleles appear to have been selectively retained in populations and the McDonald-Kreitman neutrality index indicates the locus is under strong positive selection. This is in contrast to genes encoding HLA which are under moderate purifying pressure (10).

The diversification of *TNFRSF13B* across species, the high frequency of dominant negative variants and the evidence of positive selection suggested to us that the biological impact of *TNFRSF13B* functions are incompletely understood. To explore potential functions of *TNFRSF13B* and pressures for diversification, we tested the impact of allele variants embodying the range of functions of *TNFRSF13B* on resistance to and transmission of *Citrobacter rodentium (C. rodentium*), in mice which models enterohemorrhagic *Escherichia coli* (*E. coli*) in humans (11). We wondered whether the high frequency of *TNFRSF13B* polymorphisms and the frequent dominant negative phenotypes across species could reflect an adaptation to resist common enteric pathogens.

## Results

### *Tnfrsf13b* controls resistance to *C. rodentium*

We first asked whether or to which extent *tnfrsf13b* mutations modify susceptibility of naïve mice to infection with *C. rodentium.* Accordingly, the smallest number of *C. rodentium* (10^8^) that reliably generates disease were given to wild type (C57BL/6 mice and to mice of the same background (i) with mono-allelic or bi-allelic *tnfrsf13b* mutations encoding mA144E (homologous to the frequent human A181E mutation) and to mice (ii) with fully disrupted *tnfrsf13b* and monitored excretion of viable organisms in stool during the ensuing month. Because gut microbiota potentially influence the virulence of *C. rodentium* (12), mutant and wild type mice used in these experiments were co-housed for 4 weeks prior to infection to allow admixture of flora.

After introduction of *C. rodentium,* wild type mice typically develop a mild diarrheal disease commencing at 5-7 days, reaching greatest severity at 7-10 days, and resolving at 18-28 days; infection typically elicits immunity that imparts enduring resistance to subsequent infection *(11).* Consistent with that experience, wild type mice exhibited maximum excretion of 10^7^ - 10^9^ viable *C. rodentium* between 7 days and 21 days post-infection and resolution by 35 days. In notable contrast, mice with mono-allelic mutations encoding A144E variants excreted on average 10-fold less viable *C. rodentium* than wild type mice (p=0.0246) and all mice with bi-allelic mutations and all but one with full disruption of *tnfrsf13b* excreted no viable organisms (p<0.0001) (Figure 1A). Perhaps more important, the total number of viable *C. rodentium* excreted during the course of the experiments by mutant mice was profoundly lower than the total number excreted by wild type mice (p=0.0002) (Figure 1B). Thus, mutations of *tnfrsf13b* that disrupt the function of the encoded protein decrease susceptibility to *C. rodentium* infection and this decrease is appreciated days before adaptive immunity might begin to be manifest.

**Fig. 1.**
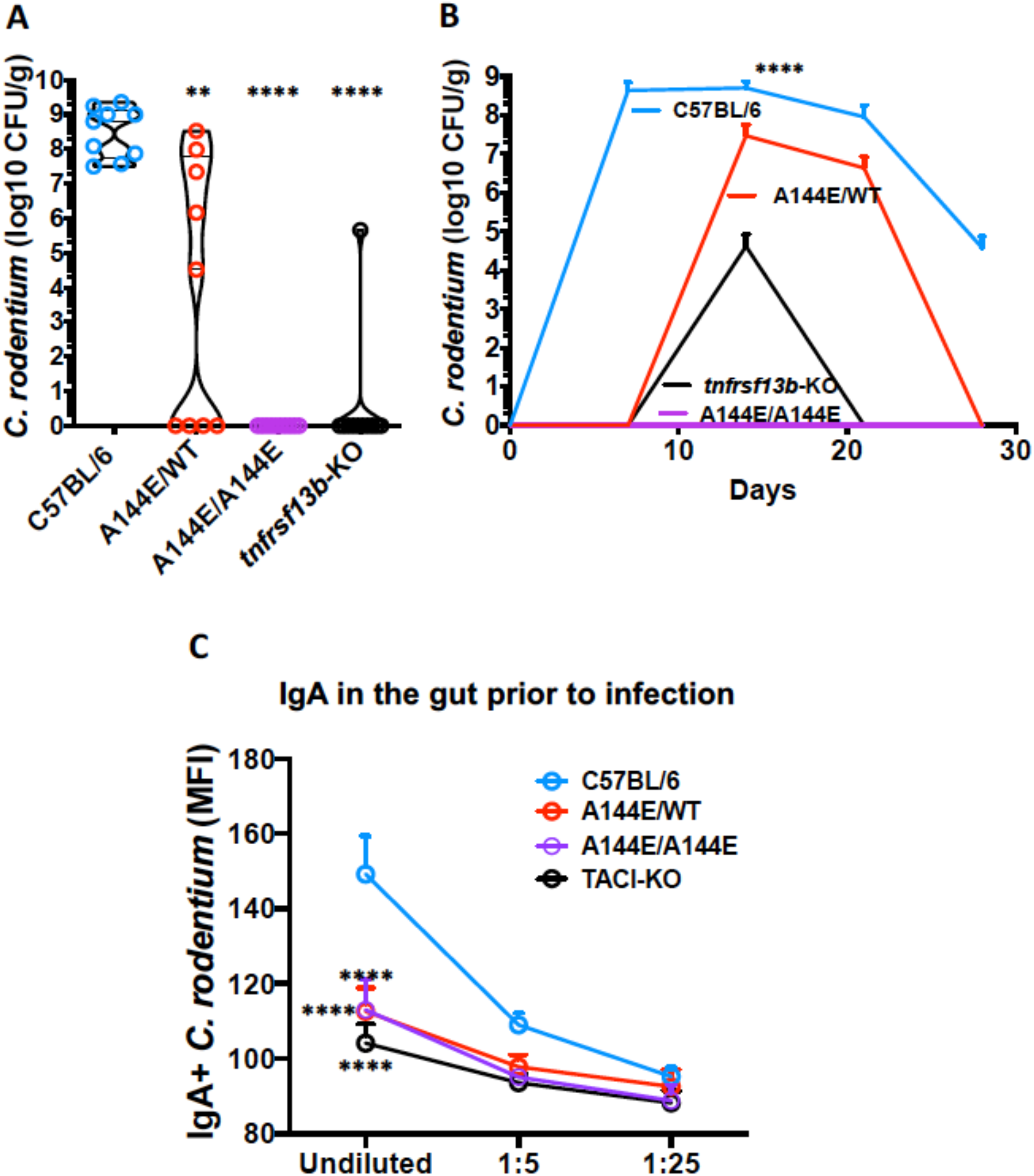
*Tnfrsf13b*-mutant mice resist infection with *C. rodentium.* To evaluate baseline resistance to enteric infection with *C. rodentium*, wild type (WT) C57BL/6 mice and mice with monoallelic or biallelic mutations in *tnfrsf13b* (A144E) and mice with targeted disruption of *tnfrsf13b* (*tnfrsf13b*-KO) were infected with 10^8^ *C. rodentium* by oral gavage and the numbers of viable organisms in stool were measured by counting CFU/g of feces after 18 hours of incubation on MacConkey plates at the peak of infection. (A) Graph depicts the maximum CFU/g of feces in the course of infection for each of the infected mouse strains. Values obtained in *tnfrsf13b*-mutant mice were compared to those in wild type by the Mann-Whitney test showing p=0.0017 for A144E/WT, p<0.0001 for A144E/A144E and for *tnfrsf13b*-KO mice. (B) Mean number of viable *C. rodentium* in stool (expressed in log10 CFU/g of feces) at various times after infection. (C) Quantification of *C. rodentium*-binding IgA in feces of mice prior to infection. IgA in the feces was measured by flow cytometry analysis of IgA+ GFP+ *C. rodentium* and detected with anti-IgA PE-labelled. Y-axis, Average of 3 independent measurements of IgA mean fluorescence intensity (MFI); X-axis, feces supernatant dilutions.

Since *TNFRSF13B* mutations confer baseline resistance to *C. rodentium*, we wondered resistance could be mediated by natural antibodies, which are known to confer resistance to other bacteria (13). Although *TNFRSF13B* mutations decrease overall production of natural antibodies (14), it is possible that natural antibodies specific for *C. rodentium* might nonetheless be present and underlie resistance in mutant mice. Indeed, *Tnfrsf13b* mutant mice had less natural IgM but no less natural IgG in the blood (Figure S1A). However, analysis of serum revealed mutant mice had no appreciable natural IgM or natural IgG in blood that could bind *C. rodentium* (Figure S1B). More important, analysis of stool from un-manipulated mutant and wild type mice revealed that wild type but not mutant mice had natural enteric IgA that bound *C. rodentium* (Figure 1C). Thus, the baseline resistance of mutant mice to *C. rodentium* infection is not mediated by natural antibodies.

If natural antibodies do not increase baseline resistance to *C. rodentium*, perhaps hastened primary antibody responses do so. To detect an early primary Ig response, we assayed the blood and stool of the various strains for presence of *C. rodentium* specific Ig 7 days after infection with that organism (Figure S2). Six of eight wild type and only 2 of 30 *tnfrsf13b* mutant mice had IgM specific for *C. rodentium* in blood seven days after infection (Figure S2A). Both wild type and *tnfrsf13b* mutant mice had *C. rodentium*-specific IgG, but the concentrations were low (<10 μg/ml) (Figure S2B). Twenty one days after infection, when adaptive IgG responses are generally detected, all wild type but fewer than one half of mutant mice had *C. rodentium* specific IgG. Since antibodies that protect against *C. rodentium* target intimin (a virulence factor) we measured the concentration of these antibodies in the serum of mice before and after infection. Intimin-specific antibodies were not observed in any strain until 14-21 days after infection and levels in wild type mice exceeded levels in mutant mice (p<0.01), (Figure S2D). Wild type mice also produced significantly more intimin-specific IgA than *tnfrsf13b*-mutant mice 21 days after infection (Figure S2E). Thus, hastened primary antibody responses do not explain heightened resistance of *tnfrsf13b* mutant mice to *C. rodentium*.

### The *tnfrsf13b* genotype determines development of virulence by *C. rodentium*

Since productive infection with *C. rodentium* requires acquisition of virulence in the host environment (12), we wondered whether mutations of *tnfrsf13b* could block such acquisition. To answer that question we assayed development of virulence by *C. rodentium* engineered to express a *ler* bioluminescent reporter gene by fusion of the *ler* promoter with the luxCDABE operon of *Photorhabdus luminescens* (*ler-lux C. rodentium*) (12). Thus, *ler-lux C. rodentium* were introduced into *tnrsf13b* mutant and wild type mice and 5 days later *ler* expression was assayed in the intestinal walls. As Figure 2 A and B show, expression of *ler* was ~8-40-fold higher in organisms introduced into wild type than into mono-allelic or bi-allelic A144E mutant mice. Decreased expression of *ler* expression in *tnrsf13b* mutant mice was paralleled by decreased excretion of viable organisms. Thus, mono-allelic and bi-allelic A144E mutant mice excreted 11-fold and 114-fold fewer viable organisms respectively than wild type mice (Figure 2C). Thus, A144E heterozygosity and the dominant negative phenotype are associated with and possibly cause nearly full suppression of *ler* expression and a profound decrease in excretion of viable *C. rodentium*. To confirm that concept, we tested whether induction of *ler* before infection vitiate differences between mutant and wild type. Figure 2D shows that inducing virulence in *C. rodentium* prior to infection by culture in DMEM medium at 37°C (15) abrogates resistance of *tnfrsf13b-*mutant mice to that organism. These results demonstrate that resistance to *C. rodentium* infection is likely exerted prior to or during acquisition of virulence.

**Fig. 2.**
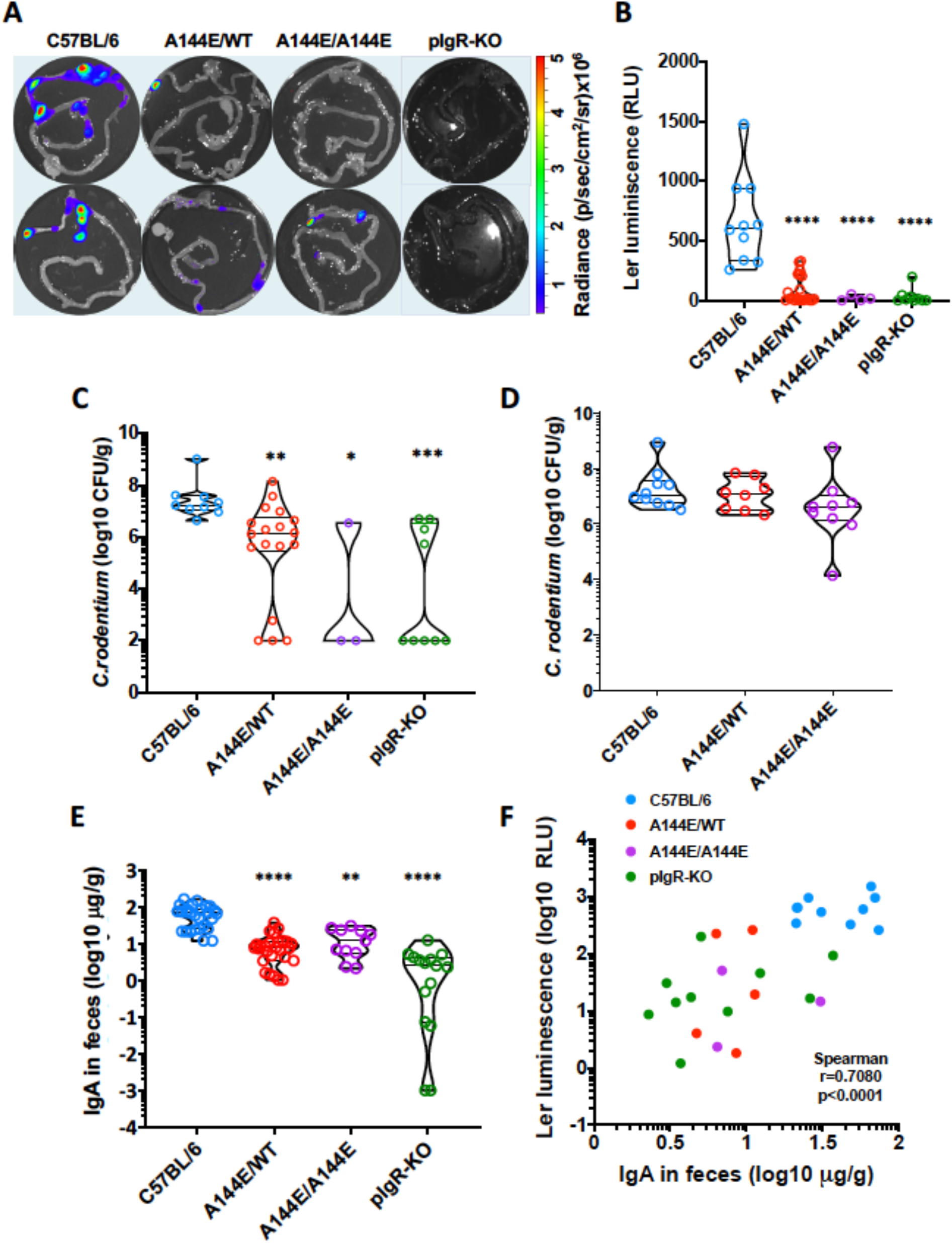
*Tnfrsf13b*-mutations block *C. rodentium* virulence gene induction. (A and B) *Ler* expression measured by bioluminescence of *ler/lux-C. rodentium* attached to the intestinal wall of mice infected for 7 days. (Graph depicts *ler* expression in relative light units, RLU, in each strain (Y-axis). RLU reflects the photons/s measured in each image divided by the background luminescence. Comparisons done with One-way ANOVA yielded p<0.0001, followed by multiple comparisons to C57BL/6 mice (control), yielded for A144E/WT, p<0.0001; A144E/A144E, p<0.0001; and for pIgR-KO, p<0.0001. (C) Graph depicts the *C. rodentium* CFU/g of feces obtained 7 days after infection for each of the infected mouse strains. Values obtained in *tnfrsf13b*-mutant mice were compared to those in wild type by the One-way ANOVA (p<0.0001) followed by the Kruskal-Wallis test showing p=0.0090 for A144E/WT, p=0.0153 for A144E/A144E and p=0.0008 for *pIgR*-KO mice. (D) Graph shows CFU/g of feces in C57BL/6 or *tnfrsf13b*-mutant mice 14 days following infection with 10^8^ *C. rodentium* grown overnight in DMEM medium and expressing *ler*. Statistical analysis was by the Kruskal-Wallis test. High virulence *C. rodentium* infects wild type and *tnfrsf13b*-mutant mice equally (p>0.05). (E) IgA concentration measured in feces 7 days after infection, by ELISA. Comparisons done with One-way ANOVA yielded p<0.0001, followed by multiple comparisons to C57BL/6 mice results (control) by the Kruskal-Wallis test, yielding for A144E/WT, p<0.0001; A144E/A144E, p=0.0049; and for pIgR-KO, p<0.0001. (F) Correlation analysis between luminescence reflecting *ler* expression by bacteria attached to the gut walls and IgA concentration in feces supernatants obtained 7 days after infection. Spearman test r=0.7080, p<0.0001 indicating that IgA concentration in feces is correlated with *ler* expression.

### Natural IgA induces *C. rodentium* virulence gene expression

Since *tnfrsf13* governs B cell maturation and production of natural and elicited antibodies we wondered whether antibodies might exert a heretofore unrecognized impact on *C. rodentium* virulence. *TNFRSF13B* mutations in humans (5, 16, 17) and *tnrsf13b* mutations in mice (3, 18) are associated with decreased production of IgA and IgA maintains homeostasis of commensal bacteria in the gut, which in turn protects against certain gut pathogens (19). However, IgA is not required for the clearance of *C. rodentium* (20). Whether natural IgA could influence *C. rodentium* virulence has not been explored. To address that possibility, we first measured IgA in feces of un-manipulated *tnrsf13B* mutant and wild type mice. Mono and biallelic A144E mutant mice had 10-100-fold less IgA in feces than wild type (Figure 2E). We next asked whether the amount of IgA in feces might influence the development of virulence by *C. rodentium*. Seven days after infection with *ler lux C. rodentium* the amount of IgA per g of feces correlated significantly with *ler* expression by organisms attached to the intestinal walls (Figures 2E, 2F and S3B) and with *C. rodentium* CFU (Figure S3A). These results were consistent with the possibility that IgA influences the acquisition of virulence by *C. rodentium*.

To determine that IgA rather than other property of *tnfrsf13b* mutant mice influences the acquisition of virulence by *C. rodentium*, we tested whether mice with wild type *tnfrsf13b* but harboring a mutation that compromises secretion of IgA into the small intestine would resist *C. rodentium* like *tnfrsf13b*-mutants. Mice lacking the polymeric Ig receptor (pIgR-KO), which transports polymeric IgA from basolateral to apical membrane of intestinal epithelium and hence have little or no IgA in the gut (21) were infected with ler lux *C. rodentium* and *ler* expression and *C. rodentium* CFU in stool were measured days later. As expected, the pIgR-KO mice had 10-10,000 fold less IgA in the feces than wild type mice at baseline. After infection with *ler lux C. rodentium*, pIgR-KO mice had on average 20-fold less *ler* luminescence (Figures 2A and 2B) and ~100-fold less *C. rodentium* CFU in the feces (Figures 2C and S3A) than wild type mice. Thus, absence of IgA rather than other facets of the *tnfrsf13b*-mutant phenotype is associated with resistance to *C. rodentium* and presence of IgA with susceptibility to *C. rodentium* infection.

Given the association of IgA in gut with susceptibility to *C. rodentium* infection, we wondered whether IgA might in some way promote virulence. To test whether IgA can promote virulence, we tested whether supernatant of feces containing or lacking IgA induce virulence in *C. rodentium.* Thus, serial dilutions of supernatant obtained from fresh feces from naive wild type, *tnfrsf13b*-mutant or PIgR-KO mice were added *C rodentium* that had been grown in LB medium under conditions that support non-virulence and *ler* expression was then assayed. As figures 3A-3C show, supernatant from feces of wild type mice induced higher (>2.5-fold) *ler* expression than supernatant from feces of *tnfrsf13b*-mutant or from PIgR-KO mice. Further, supernatant from wild type feces administered by gavage to *tnfrsf13b*-mutant mice restored virulence (Figure 3D). Furthermore, *ler* expression directly correlated with concentration of IgA (Figure 3E), consistent with IgA inducing virulence. To test whether IgA (the predominant Ig in stool) or IgG can directly induce virulence, purified polyclonal IgA (Figures 3E-F) and IgG (Figure S4A) were added to cultures of *ler lux C. rodentium* and luminescence was assayed 1 hour later. As Figure 3 and Figure S4A show, both IgA and IgG induced *ler* expression by *C. rodentium* in a concentration dependent manner. However since IgG is not present in the gut early during infection it is unlikely that “natural” IgG contributes to the induction of virulence in a non-inflamed gut. The impact of IgA/IgG on *C. rodentium* virulence did not depend on complement as heating the supernatants to 56°C for 30 minutes to inactivate complement (22) did not efface induction of *ler* expression (Figure S4). Finally, we asked whether virulence-inducing IgA necessarily reflected production by mutant mice or could have been acquired from maternal transfer. Assay of IgA in stool of various combinations of mutants or wild type progeny of mutant or wild type mothers revealed the concentration of IgA in the gut was determined by the genotype of the progeny and not by the mother s’ genotype (Figure 3G).

**Fig. 3.**
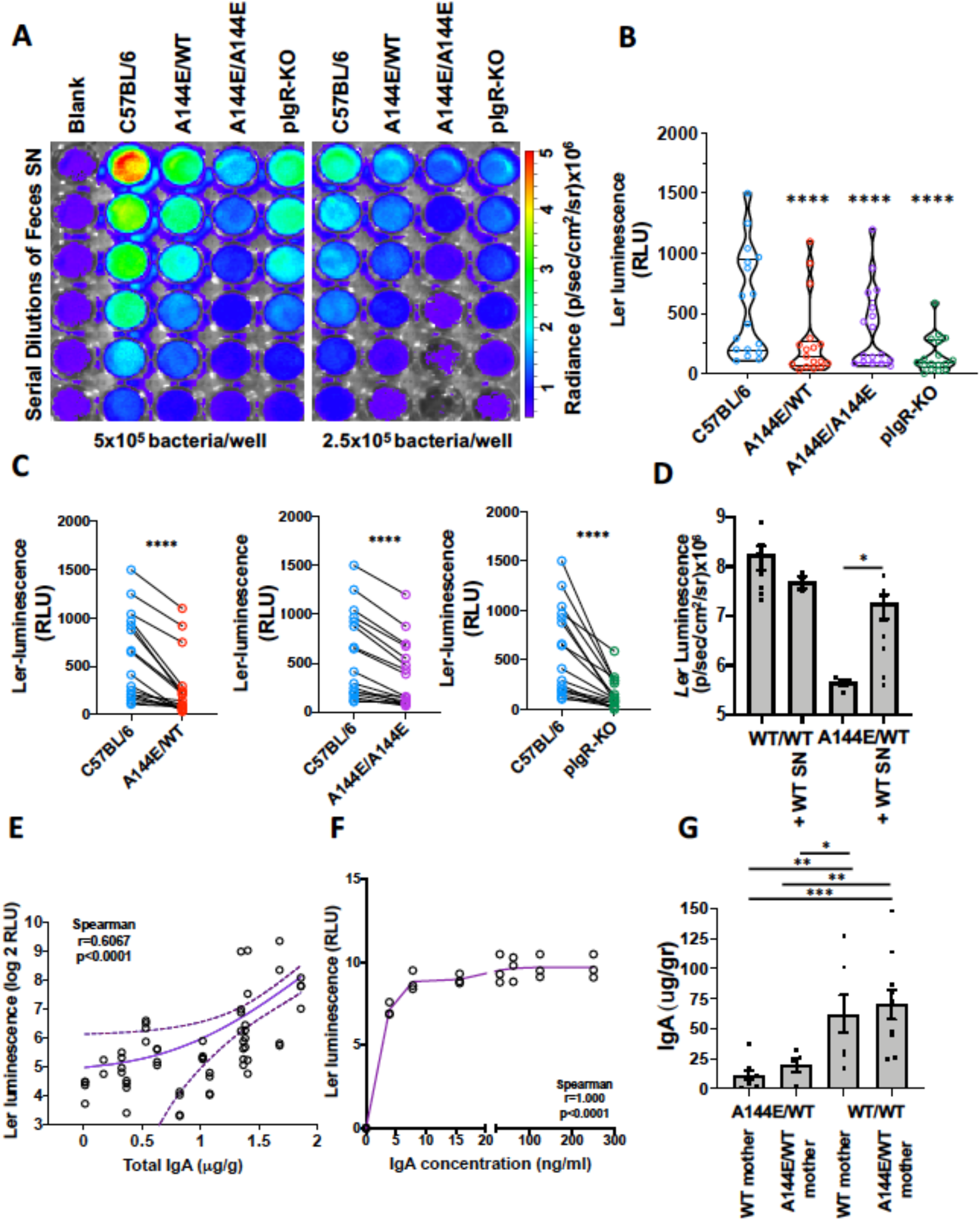
*Ler* expression increases directly with IgA concentration. *Ler* expression measured by bioluminescence imaging (BLI) of *ler/lux C. rodentium* incubated with serial dilutions of feces supernatants obtained from non-infected naïve C57BL/6, A144E/WT, A144E/A144E or pIgR-KO mice for one hour at 37°C. (A) Example of a plate reading in a typical experiment. (B) Graph shows *ler* expression detected with BLI and expressed in Relative Luminescence Units (Y-axis), normalized for the feces weight in 12 independent experiments for each mouse strain (X-axis). Luminescence data in each mutant mouse was compared to the luminescence in the C57BL/6 group by One-way ANOVA (p<0.0001) followed by the Holm-Sidak’s multiple comparisons test (p<0.0001, ****). (C) Paired analysis of normalized (to the feces weight) individual measurements of *ler* expression (Y-axis), comparing luminescence obtained with supernatant (SN) from mutant mice with that from C57BL/6 mice, within each experiment (X-axis). Paired t test analysis yielded p<0.0001, ****. (D) *Ler* expression measured by bioluminescence of *ler/lux-C. rodentium* attached to the intestinal wall of WT or A144E/WT mice infected for 7 days and treated (or not) with feces SN obtained from naïve WT mice. Feces SN (200μl, undiluted) were administered by gavage twice, one day prior and 2 days after infection. Gavage of WT SN increased *C. rodentium* virulence following infection of A144E/WT mice. Comparisons done with Mann-Whitney tests yielded p= 0.0212. (E) Regression and correlation analysis between IgA concentration in the feces SN (X-axis) and *ler* expression (Y-axis). Continuous line represents the average and dotted lines the 95% confidence limit. The slope of the curve was different from 0 with p<0.0001. Analysis by the Spearman test yielded r =0.6067 and an approximate p<0.0001 (two-tailed). (F) Ler-lux *C. rodentium* was incubated with serial dilutions of isolated polyclonal murine IgA in PBS for one hour at 37°C. Y-axis, *ler* expression detected with bioluminescence imaging (BLI), X-axis, IgA concentration in ng/ml. (G) IgA concentration in feces of A144E/WT of WT mice born from WT or A144E/WT mothers. Mann-Whitney tests yielded p<0.001, ***; p<0.01, **, p<0.05, *.

### The Ig genes encoding *C. rodentium-*bound enteric IgA+ in C57BL/6 and in A144E *tnrsf13B*-mutant mice have distinct properties

Our findings suggest that *C. rodentium* apparently evolved to co-opt some property of gut IgA to signal induction of the virulence program and various disruptive mutations of *tnfrsf13b* in mice and *TNFRSF13B* in humans were sustained possibly to avert infection and transmission of this class of organisms. Since IgG, which is not present to any great extent in normal gut, and Ig from all wild type animals yet tested induces virulence in *C. rodentium*, we reasoned that the “active” region of Ig likely resides in F(ab)2 domains. To test that concept, we compared the sequences of IgH and IgL genes from IgA+ *C rodentium*-specific B cells isolated from the Peyers patches of mice 14 days after infection or 5 days after re-infection. Following re-infection, of 10 clones isolated from wild type mice, two, B4 IgC03 and D3 IgC09, were found repeatedly (3 and 2 times, respectively). All of the clones were encoded by one of 3 VH regions VH1, VH3 or VH5. Six of ten HC sequences were >98% homologous to germline (Tables S1 and S2). The HC Ig sequences also had short CDR3 and short N-regions (non-templated nucleotide additions during V(D)J recombination), as would be expected of antibodies arising from fetal lineage precursor cells with restricted terminal deoxynucleotidyl transferase (TdT) activity (23). Only 59% of the mutations observed were non-synonymous suggesting absent or weak antigen selection (Table S2). These properties are characteristic of natural antibodies (24). In contrast, the sequences of IgH from A144E homozygous mice were characteristic of elicited responses. Upon re-infection, only 40% of the clones from mutant mice contained germline sequences and none were repeated, indicating greater diversity than the wild type (Tables S3 and S4). *C. rodentium*-specific IgA clones from A144E homozygous mice had longer N regions than clones from wild type mice and although the overall mutation frequency in clones from A144E homozygous mice was low, on average (3.6%), 70% of the mutations were non-synonymous suggesting weak antigen selection. *C. rodentium*-specific IgA clones from A144E homozygous mice had longer heavy chain CDR3s (on average the HC CDR3s from A144E homozygous mice were 3 amino acids longer than the HC CDR3s from wild type mice) (Figures 4A-C). *Tnfrsf13b*-mutant mice did not have decreased frequency of *C. rodentium*+ IgA B cells or of CD19+ B cells in the Peyers patches (Figure S5B) compared to WT mice; *Tnfrsf13b*-mutant mice had as many T cells and increased frequency of B cells in the spleen compared to WT mice (Figure S5C).

**Fig. 4:**
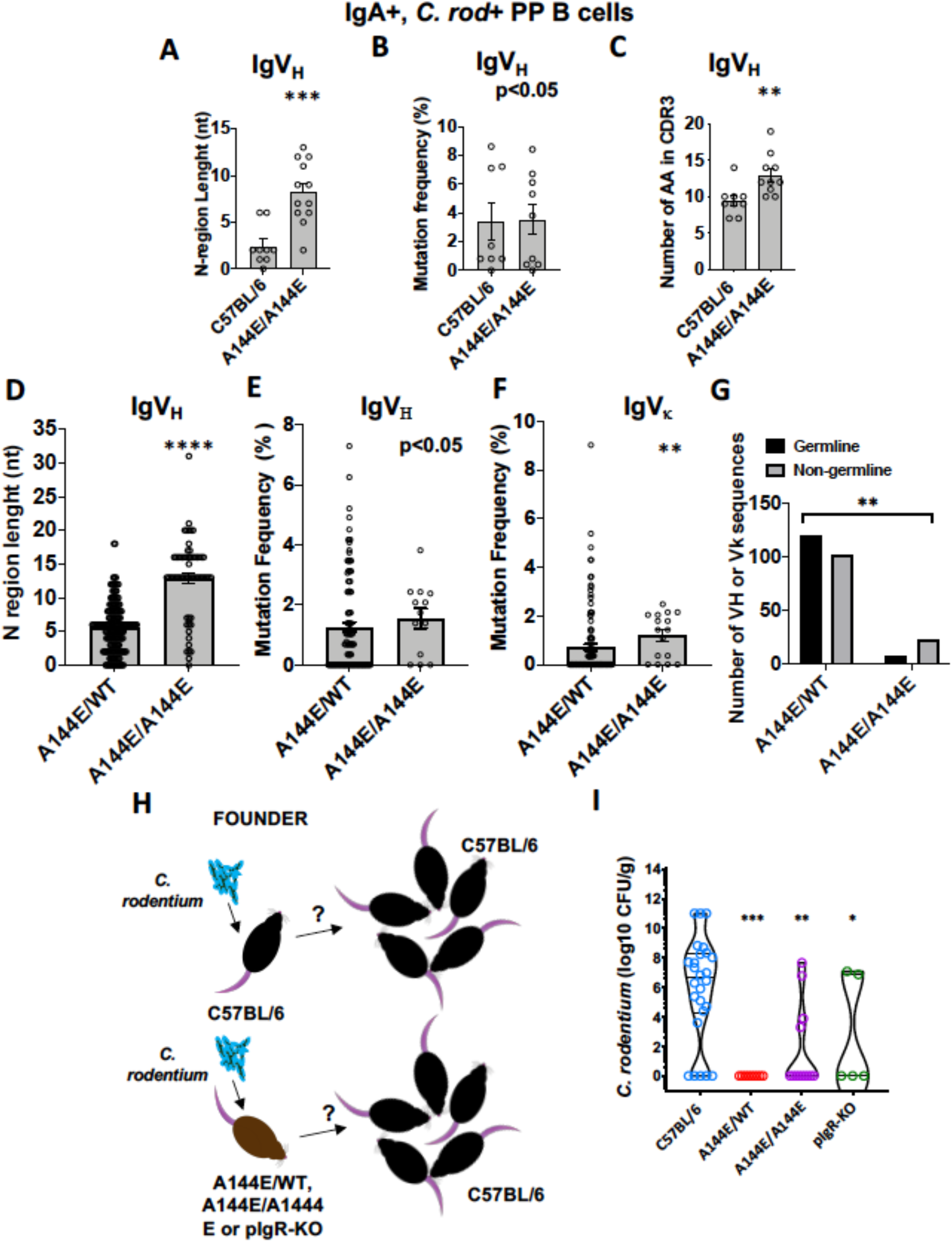
IgA sequences from sorted *C. rodentium*-bound IgA+ B cells of C57BL/6 and *tnfrsf13b*-mutant mice have distinct properties and determine *C. rodentium* spreading. Single IgA-positive GFP-*C. rodentium*-bound B cells were sorted from the Peyers patches of infected C57BL/6, A144E/WT or A144E/A144E mice either 5 days following re-infection (A, B and C) or 14 days following primary infection (D, E, F and G) (See also Figure S5 and Tables S1-S7). IgA H+L sequences were obtained from cDNA by PCR followed by Sanger sequencing (A, B and C) or by next generation sequencing of barcoded single cell barcoded cDNA libraries (D, E and F). (A and D) Graphs compare the lengths of the N-regions (number of nucleotides) of CDR3 regions (in amino acids) of HC IgA sequences obtained from C57BL/6 A144E/WT or A144E/A144E mice. (B, E and F) Graphs depict the frequencies (%) of mutated nucleotides in VH or V_H_ exons relative to their closest germline in HC sequences obtained from C57BL/6, A144E/WT or A144E/A144E mice. (C) Number of aminoacids (AA) in the complementary determining regions of Ig VH sequences obtained from C57BL/6 and A144E/A144E mice. Mann-Whitney test analysis yielded p=0.0066 (A), p=0.9196 (B), p=0.002 (C), p<0.0001 (D), p=0.1145 (E) and p=0.0036 (F). (G) Contingency analysis of frequency of germline *C. rodentium* binding IgA, VH and Vk sequences obtained from A144E/WT or A144E/A144E mice. Chi-square test yielded a p=0.016 indicating that germline sequences are rarer in A144E/A144E mice compared to A144E/WT mice. (H) Schematic of experimental protocol. Founder mice are either C57BL/6, *tnfrsf13b*-mutant mice or pIgR-KO mice. Two founder mice were co-housed with 3 C57BL/6 mice/cage. Only founders were inoculated with low virulence 10^8^ *C. rodentium.* CFUs were counted on non-primarily infected mice 7 days after infection. (I) Graph depicts CFU/g of feces in mice not primarily infected according to the strain of the inoculated founders (depicted in the X-axis). Comparisons were by Kruskal-Wallis tests followed by Dunn’s multiple comparison tests. Kruskal-Wallis yielded p<0.0001; Dunn’s multiple comparison tests yielded p= 0.0111, *; p=0.0060, **, p=0.0004***.

To determine what properties of IgA were attributable to the genotype as opposed to those determined by the environment, we sequenced IgA (H+L) from single cells using 10X GEM technology obtained from A144E/WT or A144E/A144E littermates, 14 days after infection. The results shown in figures 4D-G and Tables S5 and S6 largely confirm that the dose of the A144E allele is associated with larger N regions and decreased frequencies of germline sequences. These results show that *tnfrsf13b* mutant mice have reduced natural IgA in the gut and instead produce adaptive IgA.

### *Tnfrsf13B* A144E or PIgR-KO alleles control *C. rodentium* spreading

*C. rodentium* might exploit germline encoded natural IgA to secure a niche and prolong residence in the gut. However, this adaptation would be eclipsed if germline-encoded IgA hindered transmission of virulent organisms. To evaluate this possibility, we examined the transmissibility of *C. rodentium* from wild type and from various mutant strains of mice to uninfected susceptible wild type mice (Figures 4G and 4H). Wild type or *tnfrsf13B*-mutant mice were infected with *C. rodentium* (founder mice) and co-housed mice were tested for infection 7 days later. Wild type mice transmitted disease to 21 of 25 co-housed wild type mice (84%). Mono-allelic A144E mice in contrast transmitted *C. rodentium* to no other co-housed mice whether wild type or mutant. Bi-allelic A144E transmitted *C. rodentium* to only 4 out of 13 (30.8%) and PIgR-KO transmitted infection to 2 of 5 (40%) co-housed wild type mice. Thus, natural IgA not only induces virulence, it also enables transmission of *C. rodentium* within a colony. Our results suggest the extraordinary frequency *tnfrsf13b* mutations might in part have been preserved to counter this vulnerability.

## Discussion

Here we report what may be the truest example of balancing selection and co-evolution. Balancing selection ideally requires: (a) variant alleles where each variant is maintained at a particular equilibrium frequency; (b) a distinct benefit afforded by some fraction of the variants and/or heterozygous advantage; (c) a biological cost. In one famous example of balancing selection, the sickle cell β–globin gene is maintained in populations because the heterozygous state affords resistance to malaria at the cost of homozygote associated sickle cell disease (25). We show in this report that *tnfrsf13b-*mutant alleles that impair receptor function enhance resistance to *C. rodentium* and limit bacterial dissemination because of decreased natural IgA which *C. rodentium* has co-opted to express virulence.

Like the sickle cell β-globin gene the advantage given by *TNFRSF13B* mutants comes with a biological cost. Indeed, *TNFRSF13B* mutants are associated with common variable immune-deficiency (CVID) (5) and auto-immunity (26). In contrast to the sickle cell β–globin gene *TNFRSF13B* polymorphisms govern many facets of immunity. We propose that *TNFRSF13B* polymorphisms, define a continuum of immune responses by controlling aspects of both innate and adaptive immunity. We (1, 2) and others (27) showed that *tnfrsf13b* governs differentiation of plasma cells, controls the synthesis of “natural IgA antibodies” [in this communication] and determines affinity maturation of antibodies by controlling functions of B and T helper follicular cells in the germinal center (4). While the sickle cell β-globin gene impacts only the carrier, *tnfrsf13b* polymorphisms might impact on the health of the community by limiting transmission of bacteria regulated by *LEE*-like loci. Our findings thus suggest that the high frequency of dominant-negative *TNFRSF13B* variants may be maintained by balancing selection.

Commensal organisms compete favorably against and hence suppress *C. rodentium* late in the course of infection (28). However, gut commensals are also required for *C. rodentium* colonization of the mucosa (29). We found the composition of the microbiota to significantly differ between *tnfrsf13b* mutant and wild type mice (Figures S6 and S7) but that in itself did not explain the relative resistance to infection by *tnfrsf13b* mutant mice given the arguments that follow: (i) Normalization of the microbiota by co-housing (for 4 weeks) did not render *tnfrsf13b* mutant mice susceptible or wild type mice resistant to *C. rodentium* infection; (ii) Transfer of feces SN from WT mice, that do not contain bacteria, to resistant *Tnfrsf13b*-mutant mice induces susceptibility; (iii) Progeny of WT or *tnfrsf13b*-mutant mice manifest the susceptibility associated with their genotype (not the susceptibility associated with their mothers’). Thus, A144E heterozygous mice, obtained by crossing wild type mice with homozygous *tnfrsf13b* A144E mice, frequently resist infection or produce 20-fold less CFUs than wild type littermates and *tnfrsf13b* A144E/A144E and *tnfrsf13b*-KO mice, produced and backcrossed to C57BL/6 independently, are resistant. These facts support the resistant phenotype is a direct consequence of *tnfrsf13b* genotype.

The prevailing concept is that IgA is protective (19, 30, 31). IgA is thought to protect against pathogens by exerting direct effects such as neutralizing virulence (decreasing motility by binding to adhesins, pili, flagella) (32–34), by promoting entrapment in the mucus layer (35), immune exclusion (36), clearance by agglutination and enchained growth (9, 37) or by direct modulation of gene expression (38, 39). In one example, natural IgA protects against *Salmonella thyphimurium* and against necrotizing enterocolitis in newborn infants (40, 41). IgA may also protect against pathogens by indirect actions such as enhancing colonization by protective (anti-inflammatory) commensal species (42). For example, *Bacteroides fragilis* capsule induces specific IgA which in turn, increases its adherence to intestinal epithelial cells (31), and commensals such as *Bacteroides thetaiotaomicron* utilize IgA to support mutualism(39). Our data instead indicate that *tnfrsf13b* mutants induce resistance to *C. rodentium* because of a relative depletion of IgA “natural” antibodies in the gut which induce *C. rodentium* virulence. Accordingly, WT feces SN (and polyclonal IgA alone) induce virulence, in accord to their IgA concentration. Furthermore, pIgR-deficient mice that have hardly any IgA in feces owing to an independent mechanism, are resistant to *C. rodentium*. Other than our report, pathogens’ adaptation to “natural” IgA to increase virulence and transmissibility has not to our knowledge been described. Because *tnfrsf13b* mutants maintain the ability to make “adaptive” IgA it is possible that protective functions of IgA are maintained.

For all the advantages IgA provides, IgA deficiency is the most common immune-deficiency with frequencies varying between 1:143 in the Arabian peninsula to 1 in 500 Caucasian individuals (43, 44). This number may be underestimated since IgA-deficiency is often asymptomatic. One of us (Dr. Raif Geha) (5, 45) showed that all of the patients with *TNFRSF13B* mutations and common variable immunodeficiency examined also had IgA deficiency and that one IgA-deficient patient also had a *TNFRSF13B* mutation. In one recent study, Pulvirenti et al. (46) showed that 13% IgA deficient patients carried at least one mutated *TNFRSF13B* allele. Most IgA-deficient individuals are asymptomatic and only a small percentage develop recurrent sino-pulmonary infections and/or auto-immune manifestations (43). Given the limited morbidity of IgA deficiency, one might wonder whether benefits conferred by constraint on virulence of *LEE*-dependent enterobacteria outweigh the detrimental impact of decreased IgA in the gut. Data showing high frequency of H and L mutations in *C. rodentium* specific B cells in mutant mice suggest enhanced (compensatory) adaptive antibody responses. In accord, IgA-deficient patients were found to have enhanced adaptive antibody responses to pneumococcal vaccination (47). Perhaps it is this type of response that explains the mild phenotype of many individuals with IgA deficiency.

We show here that a common *tnfrsf13b* variant even when expressed from a single allele together with the wild type gene induces resistance to an enterohemorrhagic microbe by blocking expression of virulence genes which in turn limit transmission and dissemination of disease. *C. rodentium* is a commonly used model for enteropathogenic infections in humans, dependent on the *LEE* locus. Given the pronounced impact of mono-allelic mutations, our findings identify a receptor that might be temporarily targeted to disrupt transmission of organisms in epidemics of organisms regulated by *LEE*-type loci.

## Materials and Methods

### Experimental Models and Subject Details

#### Mice

C57BL/6 wild type (WT) mice were purchased from The Jackson Laboratory (C57BL/6J, The Jackson Laboratory, IMSR Cat# JAX:000664; RRID: IMSR_JAX:000664). Polymeric Ig receptor-knockout (pIgR-KO) (48), *Tnfrsf13b*- KO mice (6) and mice harboring bi-allelic (A144E/A144E) or mono-allelic A144E variants (WT/A144E) (14), *Tnfrsf13b*, homologous to the human A181E, were previously described. All the knockouts and *Tnfrsf13b*-mutant mice were bred into the C57BL/6 background. In some experiments A144E/WT and WT littermates were obtained from crosses between WT mothers and A144E/WT males or from crosses between A144E/WT mothers and WT males. Animals of both genders between 8-20 weeks of age were maintained under specific pathogen-free conditions and all the experiments were performed in accordance with the approved animal protocol and the regulations of University of Michigan Committee on the Use and Care of Animals.

#### Citrobacter rodentium

The kanamycin (Km)-resistant wild type *Citrobacter rodentium* strain DBS120 (pCRP1:Tn5) was a gift of Dr. David Schauer, Massachusetts Institute of Technology, Massachusetts, USA (49); *C. rodentium* expressing GFP (50) used to identify *C. rodentium*-bound B cells, was a gift of Dr. Bruce Vallance, University of British Columbia, Vancouver, Canada; *C. rodentium* expressing a plasmid containing a *ler/lux* transcriptional fusion was previously described (12). Bacteria were grown overnight in Luria-Bertani (LB) broth supplemented with Km (50 μg/ml) with agitation at 225 *rpm*, at 37°C. To produce high and low virulence inoculum, 1 ml of bacteria suspension was added into either 9 ml of autoclaved LB with Km (low virulence) or to 9 ml of DMEM (high virulence). Those cultures were incubated in agitation at 37°C for 6 more hours, bringing both culture concentrations to 10^8^ CFU/ml.

### Methods Details

#### Infections

Mice were infected by oral gavage with 0.2 ml of PBS containing approximately 10^8^ CFU. At the designated time points, the feces were collected, weighed, suspended in 1 ml of PBS and serially diluted. Optimally diluted feces were plated on MacConkey plates and cultured at 37°C overnight. The number of colonies was counted and the number of bacteria per 1 g of feces was calculated. Infection was considered cleared when no colonies were detected in the undiluted feces suspension. In some experiments, mice were given naïve C57BL/6 feces supernatants by gavage (200 μl) twice, 24 hours before infection and 2 days after infection.

#### Measurement of *ler* expression

*Ler* expression was determined by measuring luminescence emitted by *ler/lux*-expressing bacteria in PBS suspensions, or in gut tissue *ex vivo*. Luminescence was detected with bioluminescence imaging (BLI) using an IVIS200 (Xenogen Corporation, Alameda, CA). When measuring luminescence of bacteria attached to the intestinal wall, the entire gastrointestinal tract was removed, bisected, washed with PBS and placed into the light-tight chamber of the CCD camera system of the IVIS 200 immediately. For PBS suspensions of feces and mouse IgA and IgG (Southern Biotech, Cat#0106-01; RRID: AB_2714214), 5 × 10^5^ ler/lux-expressing *C. rodentium* were incubated for 1 h at room temperature with feces supernatant serial dilutions or different concentrations of mouse IgA.

Luminescence emitted from *lux*-expressing bacteria in the tissue was quantified using the Living Image Software v.4.7.2 (IVIS Imaging Systems, Xenogen Corporation, RRID:SCR_014247). Relative Luminescence Units were obtained by dividing the total light measured in photons/s per unit of area in 10 minutes and dividing that number by the background luminescence, followed by standardization for the weight of feces when supernatants were used. Quantitative real time RT-PCR (qPCR) for *ler* was performed using a SYBR green PCR master mix to confirm that luminescence reflected *ler* expression according to Kamada et al. (12). Briefly, the expressions of *ler* and *Km resistant protein* were detected by qPCR using respectively 5’-AAT ATA CCT GAT GGT GCT CTT G-3’ and 5’-TTC TTC CAT TCA ATA ATG CTT CTT-3’, and 5’-CTG AAT GAA CTG CAG GAC GA-3’ and 5’-ATA CTT TCT CGG CAG GAG CA-3’ and the expression of the virulence factor *ler* was normalized to the expression of *Km resistant protein*. The relative expression of *ler* was determined as fold change when compared to the expression of bacteria cultured in LB medium.

#### Detection of total C. rodentium-reactive Igs

To detect mouse Ig Nunc MaxiSorp ELISA plates were coated for 1h at room temperature with goat anti-mouse Ig (H+L) (4 μg/ml; Southern Biotech, Cat#1010-01; RRID: AB_2794121). *Citrobacter rodentium*-specific Ig was quantified by coating Nunc MaxiSorp ELISA plates with 10^8^ bacteria/well heat-inactivated at 60°C for 1 h. After blocking, the plates were incubated with serial dilutions of mouse sera or feces supernatant for 1h at room temperature. Bound IgG, IgM or IgA were detected by adding HRP-conjugated goat anti-mouse IgG (4 μg/ml; Southern Biotech, Cat#1030-05; RRID: AB_2619742), goat anti-mouse IgM (4 μg/ml; Southern Biotech, Cat#1020-05; RRID: AB_2794201) or goat anti-mouse IgA (4 μg/ml; Southern Biotech, Cat#1040-05; RRID: AB_2714213). The reaction was visualized by subsequent addition of 2,2′-Azino-bis (3-ethylbenzthiazoline-6-sulfonic acid) substrate (Southern Biotech, #0202-01).

#### Flow Cytometry, Sorting and Antibodies

Peyer’s patches’ lymphocytes were isolated from naïve or infected mice as reported by Tsuji et al. and Cascalho et al. (1, 51). Cell viability was assed via BD Horizon Fixable Viability Stain 780 (FVS780; 1.11 μg/ml; BD Biosciences, #565388) and cells were stained with APC rat anti-mouse CD19 (1D3; 10 μg/ml; BD Biosciences, Cat#550992; RRID: AB_39848), PE rat anti mouse IgA (mA-6E1, 4 μg/ml; Thermo Fisher Scientific, Cat#12-4204-82; RRID: AB_465917) and *C. rodentium* expressing GFP. Staining were performed in 10^6^ cells, data were acquired with a BD FACS Canto II (BD Biosciences, Franklin Lakes, NJ) and 100,000 events were analyzed with FlowJo v.10.6.1 (FlowJo, RRID:SCR_008520). Single cell sorting was done on 96-well plates using a FACS Aria II (BD Biosciences, Franklin Lakes, NJ) in the Biomedical Research Facilities Core at the University of Michigan.

To quantify IgA binding to *C. rodentium* in non-infected mice, feces supernatants were diluted at 10 mg/ml in PBS and binding of feces IgA to *C. rodentium* was detected by flow cytometry. Briefly, 10^8^ *C. rodentium* expressing GFP were incubated with various dilutions of feces supernatant for 30 mins at 4°C. Bacteria were then washed and bound IgA was detected with a PE-conjugated rat anti-mouse IgA (mA-6E1, 4 μg/ml; Thermo Fisher Scientific, Cat#12-4204-82; RRID: AB_465917). Data were acquired with a BD FACS Canto II (BD Biosciences, Franklin Lakes, NJ), 100,000 events were recorded in the *C. rodentium*-GFP gate, and were analyzed with FlowJo v.10.6.1 (FlowJo, RRID:SCR_008520).

#### Ig gene sequencing

IgA-positive GFP-*C. rodentium*-positive B cells obtained from the Peyer’s patches of mice re-infected with *C. rodentium* 5 days earlier were single cell-sorted into of U-bottom 96-well plates according to methods adapted from Tiller et al. (52). In brief, cells were sorted into PCR plates containing 4 μl/well of ice-cold 0.5X PBS supplemented with 10 mM DTT (Invitrogen, Cat#Y00147), 8 U RNasin Ribonuclease Inhibitor (Promega, Cat#N2115), and 3 U Recombinant RNase Inhibitor (Takara, Cat#2313A), sealed and immediately frozen on dry ice. Total RNA from single-sorted B cells was reverse transcribed in the original sorting plate with 150 ng Random Hexamer Primer (Thermo Scientific, Cat#S0142), 1 mM dNTP Mix (Invitrogen, Cat#18080044), 7 mM DTT (ThermoScientific, Cat#P2325), 0.5% v/v IGEPAL CA-630 (Sigma-Aldrich, Cat#I3021-50ML), 4 U RNasin Ribonuclease Inhibitor (Promega, Cat#N2115), 6 U Recombinant RNase Inhibitor (Takara, Cat#2313A) and 50 U Superscript III reverse transcriptase (Invitrogen, Cat# 18080-044) and nuclease-free water in a final volume of 14 μl/well. Reverse transcription was performed at 42°C for 5 min, 25°C for 10 min, 50°C for 60 min and 94°C for 5 min. cDNA was stored at −20°C. Mouse *Igh*, *Igk* and *Igl* V gene transcripts were amplified by two rounds of semi-nested (*Igh*) or nested (*Igk* and *Igl*) PCR in 96-well plates containing 200 nM each primer or total primer mix (Table S7), 300 μM of dNTP Mix (Invitrogen, Cat#18080044) and 1.2 U HotStart Taq DNA polymerase (Qiagen, Cat#203205). The first round of reactions was performed with 3.5 μl of cDNA at 94°C for 15 min followed by 50 cycles of 94°C for 30 s, 56°C (*Igh*) or 50°C (*Igk*) or 58°C (*Igl*) for 30 s, 72°C for 55 s, and final incubation at 72°C for 10 min. After identification of the Light chain genes the semi-nest or nested second round PCR was performed with 3.5 μl of first round PCR product as template and combinations of V, J and C primers (Table S7) at 94°C for 15 min followed by 50 cycles of 94°C for 30 s, 60°C (*Igh*) or 45°C (*Igk*) or 58°C (*Igl*) for 30 s, 72°C for 45 s, and final incubation at 72°C for 10 min. PCR products were treated with ExoSAP-IT PCR Product Cleanup Reagent (Applied Biosytems, Cat#78201.1.ML) and sequencing was performed by the Sanger method at the University of Michigan Sequencing Core. Analysis of the sequences was done by using the IMGT portal (53–55) (RRID:SCR_011812), alignments were by Multiple sequence alignment by Log-expectation, MUSCLE software (56, 57) (RRID:SCR_011812). Comparisons of the amino acid composition of *C. rodentium*-specific IgA heavy chain (HC) was done using the Kullback-Leibler logotype using the Seq2Logo 2.0 software (58) (http://www.cbs.dtu.dk/biotools/Seq2Logo/index.php). Primer sequences are described on Table S8. Alternatively, IgA-positive GFP-*C. rodentium*-positive B cells obtained from the Peyer’s patches of mice infected with *C. rodentium* 14 days earlier were sorted as explained above in PBS 0.004% BSA and up to 10,000 cells were analyzed by Chromium Next Gel Bead-in-Emulsions (GEM) Single Cell V(D)J Technology. Briefly, cells were identified via generation of GEMs by combining barcoded Single Cell V(D)J 5’ Gel Beads v1.1, a master mix with cells (Chromium Next GEM Single Cell 5′ Library and Gel Bead Kit v1.1; 10x Genomics Cat#1000165), and partitioning Oil on Chromium Next GEM Chip G (Chromium Next GEM Chip G Single Cell Kit; 10x Genomics Cat#1000127) and reverse transcription and cDNA amplification were performed as recommended by the manufacturer. Next, the targeted enrichment from cDNA was conducted with the Chromium Single Cell V(D)J Enrichment Kit, Mouse B Cell (10x Genomics Cat#1000072). The cDNA quality control (QC) analysis was carried out in an Agilent 2100 Bioanalyzer (Agilent Technologies, Santa Clara, CA) using the Agilent High Sensitivity DNA Kit (Agilent Technologies Cat#5067-4626). The V(D)J enriched library was then constructed via Chromium Single Cell 5’ Library Construction Kit (10x Genomics Cat#1000020) and libraries were sequenced in a NovaSeq™ 6000 Sequencing System (Illumina, San Diego, CA). V(D)J sequences were collapsed using Cell Ranger: V(D)J Pipelines (10x Genomics, Pleasanton, CA, RRID:SCR_017344) and the V usage and clonotype profiles were generated and visualized by Loupe VDJ Browser. We were able to recover between 200 to >8,000 cell barcodes per sample.

#### 16S RNA sequencing

DNA was extracted from feces pellets using the Qiagen MagAttract PowerMicrobiome DNA/RNA EP kit (QIAGEN, Cat#27500-4-EP). The V4 region of the 16S rRNA-encoding gene was amplified from extracted DNA using the barcoded dual-index primers developed by Kozich et al. (59) (Table S8). Samples were amplified using AccuPrime Taq DNA Polymerase, high fidelity (Invitrogen, Cat# 12346086) at 95°C for 2 min followed by 30 cycles of 95°C for 20 s, 55°C for 15 s and 72°C for 5 min, and final incubation at 72 °C for 10 min, purified using a magnetic bead capture kit (Agencourt AMPure; Beckman Coulter, Cat#000130) and quantified using a fluorometric kit (Quant-iT PicoGreen dsDNA Assay Kit; Invitrogen, Cat#P7589). Purified amplicons were pooled in equimolar concentrations with a SequalPrep Normalization Plate Kit (Applied Biosystems, Cat#A1051001) and sequenced on Illumina MiSeq System (RRID:SCR_016379). Bioinformatic analysis was done using the Mothur v.1.42.3 software package (60) (RRID:SCR_011947) available at the University of Michigan Microbial Systems Laboratory.

### Quantification and Statistical Analysis

#### Statistics

All comparisons were done with GraphPad Prism v.8.0.0 software (GraphPad Software, RRID:SCR_002798). When an assumption of normal distributions could not be made values in more than two groups were compared using Kruskal-Wallis test followed by a Dunn’s multiple comparison tests. Comparisons of two groups were done by Mann-Whitney test or Wilcoxon test for paired analysis. When an assumption of Gaussian distribution could be made averages were compared by unpaired t test or when comparing more than 2 groups, using One-way ANOVA followed by multiple comparisons tests. Correlations were determined by the Spearman rank test. Differences in bacterial community structure were analyzed using analysis of molecular variance (AMOVA) in Mothur v.1.42.3 (60) (RRID:SCR_011947). Data are shown as mean ± SEM. A *P* value of equal or less than 0.05 was considered significant. Further information about which the statistical tests used in each experiment and experimental number can be found in the Figure Legends.

## Data and Code Availability

The published study includes all datasets analyzed during this study.

## Author Contributions

Conceptualization, M.C. and J.L.P.; Methodology, M.C. and J.L.P.; Experimental, M.G.M.B., D.H., A.R.L., J.K., C.K., C.M.B. and C.W.; Writing – Original Draft, M.C.; Writing – Review & Editing, M.C., M.G.M.B., C.M.B., C.W., R.G., R.B., G.N., N.K. and J.L.P.; Funding Acquisition, M.C. and J.L.P.; Resources, M.C., C.M.B., and J.L.P; Supervision, M.C. and J.L.P

## Acknowledgments

We would like to acknowledge Mr. Tyler Cox who helped with the drawings on figure 4. Microbiome data was produced by the University of Michigan Microbial Systems Molecular Biology Laboratory.

## Supplementary Materials

**Supplementary Text**

**Figures S1 to S7**

**Tables S1 to S9**

