## Supplemental material for "Natural IgA and *TNFRSF13B* polymorphism: a double edged sword fueling balancing selection"

**This PDF file includes:**

Supplementary Text
Figs. S1 to S7
Tables S1 to S8

**Supplementary text:**

### Data and Materials Availability

Further information and requests for resources and reagents should be directed to and will be fulfilled by the Lead Contact, Marilia Cascalho. This study did not generate new unique reagents. Data not included in the main text such as microbiota S16 sequences and IgA sequences will be made available on the Sequence Read Archive-NCBI data base.

### Natural antibodies in *Tnfrsf13b* mutant mice prior to infection

Consistent with prior reports (*3*), *tnfrsf13b*-mutant mice had decreased "natural" IgM but as much or more IgG in the blood than their wild type controls (Figure S1A).

### *Tnfrsf13b* mutant mice mount recall responses

Productive infection and/or *C. rodentium*-specific IgG titer predict the outcome of a secondary infection. Thus, mice that had a productive primary infection with peak CFU/g of feces $>10^5$ and/or developed anti-*C. rodentium* IgG titers above $10^{-4}$ during primary challenge resist secondary infection independently of the genotype (Figures S2F-G). Thus, *Tnfrsf13b* genotype governs baseline resistance and immunity to *C. rodentium.* The mechanisms of resistance and immunity are distinct. While baseline resistance to *C. rodentium* does not depend on acquired immunity; enhanced clearance upon re-infection is clearly dependent on acquired immunity.

### Microbiota composition in *Tnfrsf13b* mutant mice

To determine how *Tnfrsf13b*-mutant alleles shaped the microbiota, we performed 16S RNA sequencing on DNA extracted from stool samples derived from wild type or *Tnfrsf13b*-mutant mice before and following *C. rodentium* infection. Results depicted in figures S6 and S7 A and B show that *Tnfrsf13b*-mutant mice express a more diverse microbiota than wild type mice prior to infection and the difference in diversity disappears with infection principally due to changes in the microbiota in wild type mice. Differences in microbiota composition do not explain resistance to *C. rodentium* since resistance persists after co-housing for 4 weeks and is exhibited by A144E/WT and not by WT/WT littermates.

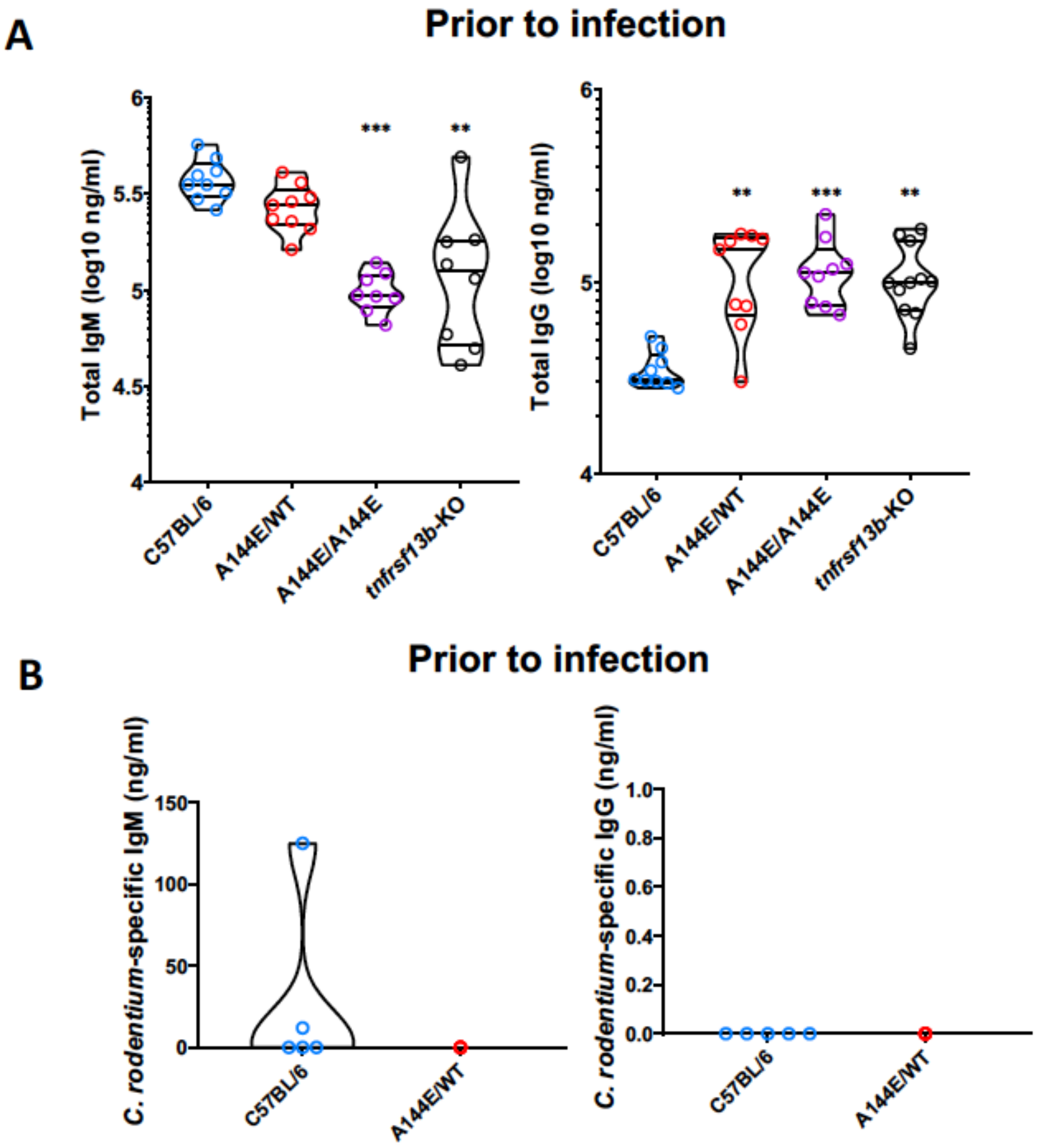


**Fig. S1. Analysis of total and *C. rodentium* or intimin-specific IgM and IgG before infection**. (A and B) Graphs represent the total IgM (left) and total IgG (right) concentrations (log10 ng/ml) in the blood before infection. Analysis was by One-way ANOVA followed by Dunnett's multiple comparisons tests comparing concentrations of IgG and IgM in *Tnfrsf13b*-mutant mice with those in C57BL/6 mice. ANOVA yielded a $p<0.0001$ (IgM) or $p=0.8198$ (IgG). Dunnett's multiple comparisons tests comparing mutants to WT yielded p=0.1121-IgM and p=0.7509-IgG, A144/WT; p<0.0001-IgM, p=0.5465-IgG, A144E/A144E; p<0.0001-IgM, p=0.3028-IgG, *tnfrsf13b*-KO. (B) Graphs represent the concentration of *C. rodentium*-specific IgM (left) or IgG (right) in blood (Y-axis) in WT or A144/WT mice prior to infection.

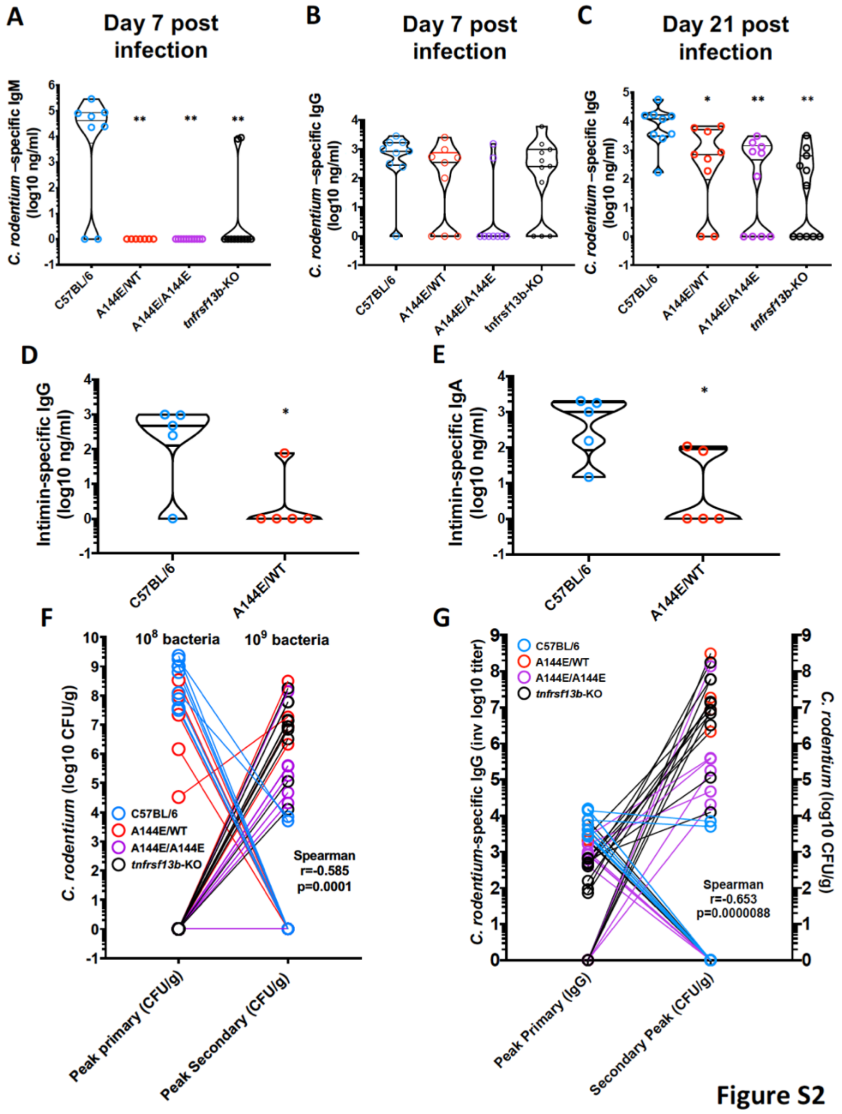


Figure S2

**Fig. S2. Primary infection and *C. rodentium*-specific antibodies protect mice from secondary infection.** C57BL/6 mice and mice with *Tnfrsf13b* A144E mutations or *Tnfrsf13b*-KO were administered $10^8$ *C. rodentium* by oral gavage and levels of antibodies against *C. rodentium* were assayed by ELISA. Anti-*C. rodentium* IgM and IgG were measured 7 days after infection; anti *C. rodentium* IgG, anti-intimin IgG or anti-intimin IgA 21 days later, a time when T cell-dependent IgG responses to *C. rodentium* achieve maximum affinity. None of the mice had detectable IgM, IgG or IgA specific for *C. rodentium* or specific for intimin in the serum prior to infection (See also Figure S1), hence the results shown reflect responses to infection. (A) Anti-*C. rodentium* IgM in serum; (B-C) *C. rodentium*-specific IgG in the blood of mice 7 days (B) or 21 days (C) after infection. Only A144E/A144E mice had significantly decreased *C. rodentium*-specific IgG. Dunnett's multiple comparisons tests yielded $p<0.01$. (D) Anti-intimin IgG in serum. (E) Anti-intimin IgA in serum. (F) Relationship between maximum CFU/g of feces in the course of primary infection with CFU/g of feces 5 days after secondary infection. (G) Relationship between anti-*C. rodentium* IgG titer in serum and CFU/g of feces of *C. rodentium* in stool 5 days after re-infection. The results indicate that having a productive primary infection or *C. rodentium*-specific IgG titer of $10^{-3}$ or below protects against a secondary challenge. Correlations were analyzed by the Spearman test and indicate that CFU/g of feces during primary infection or anti-*C. rodentium* IgG titer and CFUs after re-infection are inversely correlated.

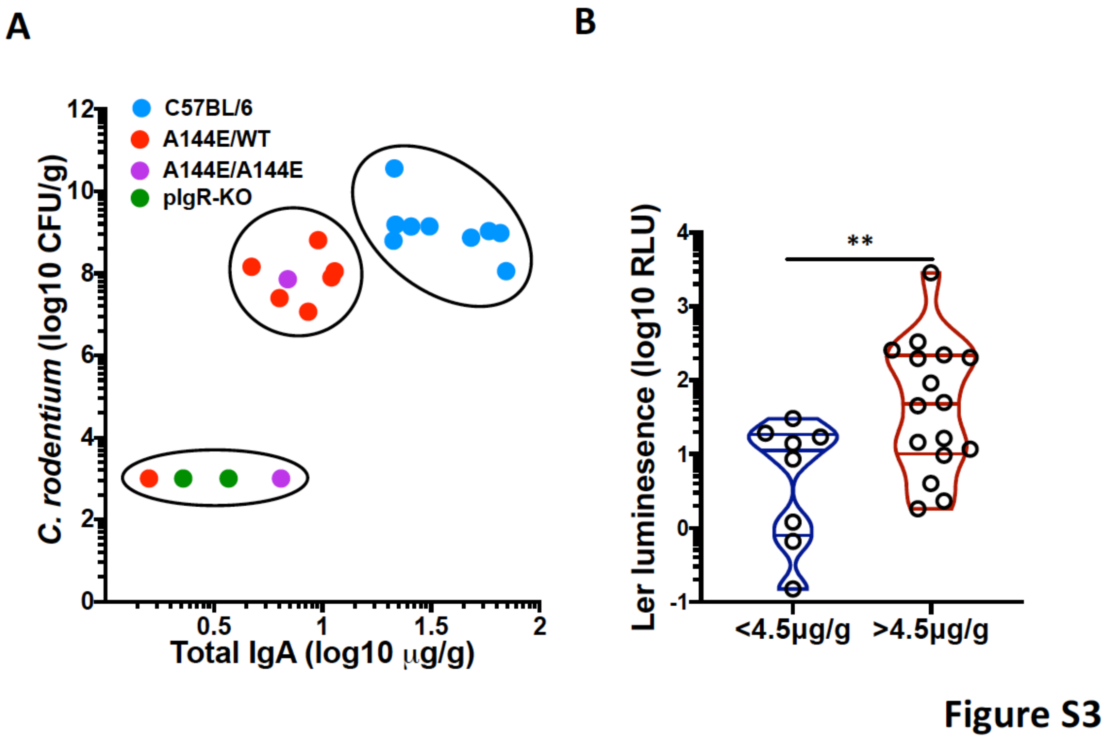


**Fig. S3**. ***Ler* expression and CFU increase with IgA concentration in the gut and IgA concentration in the gut is determined by the genotype.** (A) IgA concentration determines CFU at 7 days post-infection. The groups reflecting the IgA concentrations in relation to CFU outcomes have been noted in the figure with oval lines. (B) *Ler* expression in bacteria attached to gut walls of mice that have less than 4.5 μg of IgA per gram of feces is significantly decreased in comparison with *ler* expression in bacteria attached to gut walls of mice that have more than 4.5 μg of IgA per gram of feces. Analysis with the Mann-Whitney test yielded P=0.0029, one-tailed.

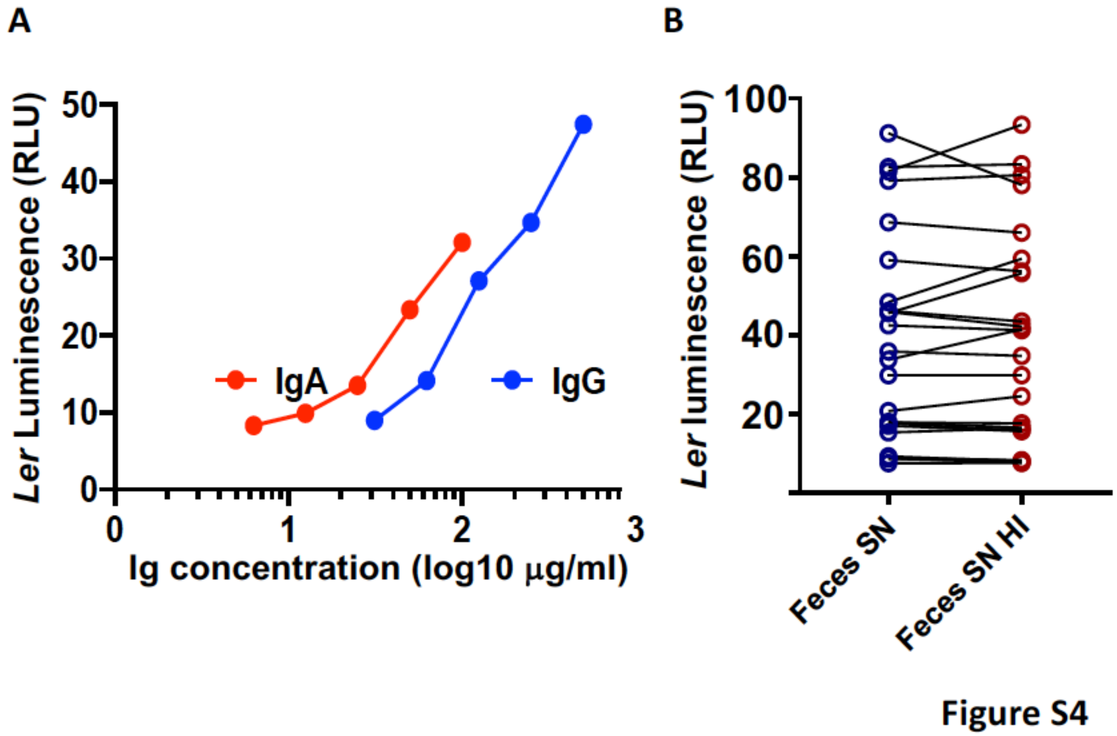


**Fig. S4: Polyvclonal IgA and IgG induce *ler* when added to *C. rodentium* and induction of *ler* is independent of heat-labile complement.** (A) Polyclonal IgG or IgA or (B) WT feces supernatant before and after heat inactivation at 56°C for 30 minutes were added to *C. rodentium* in culture. Luminescence (*ler-lux* luminescence, Y-axis) was detected with bioluminescence imaging (BLI) using an IVIS200 (Xenogen Corporation, Alameda, CA). Figure shows that heat inactivation does not inhibit virulence induction.

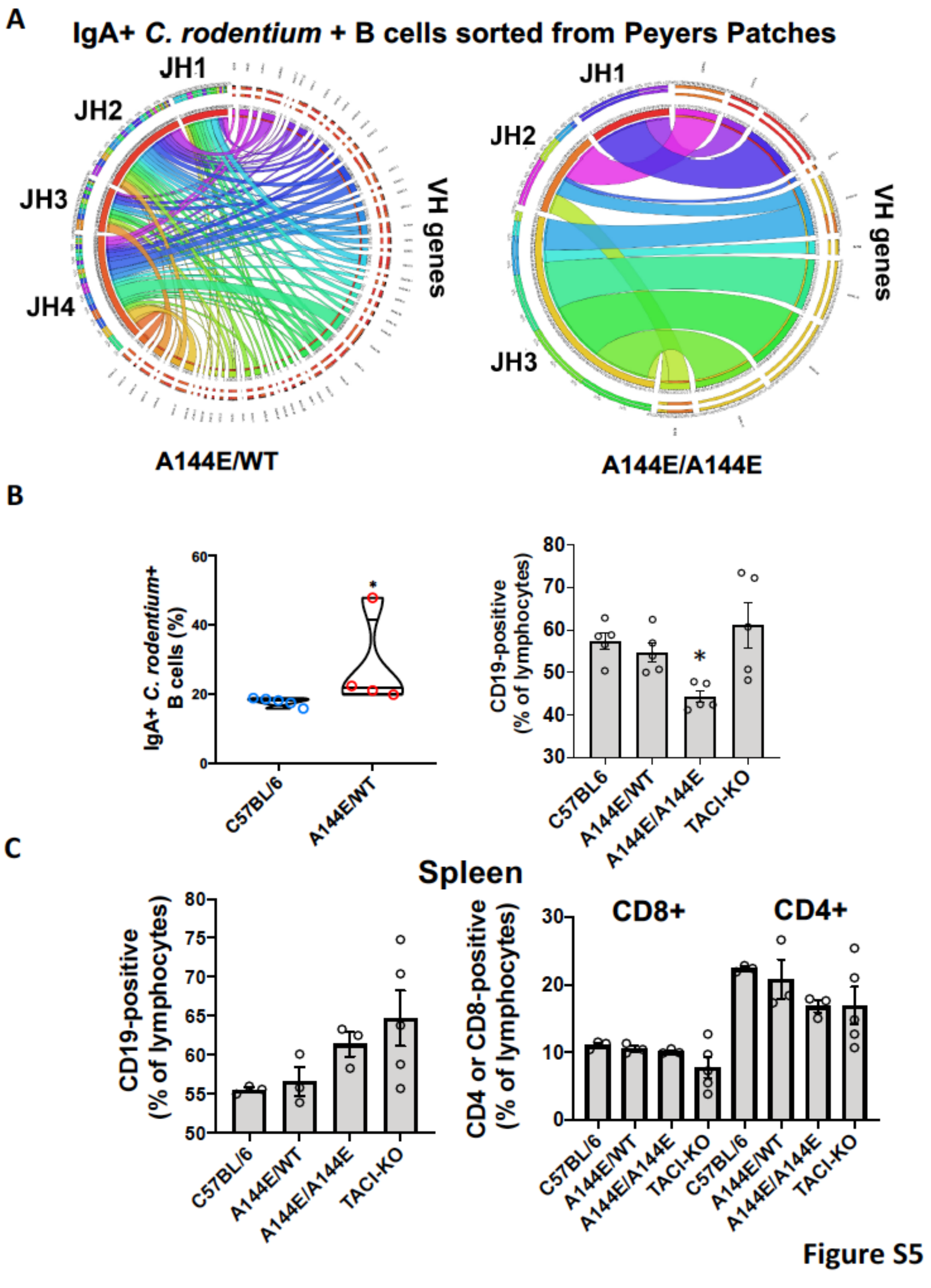


**Fig S5. Flow cytometry analysis of lymphocytes 14 days after infection and IgH repertoire diversity circos plots of Peyer's patches IgA-positive, *C. rodentium*-positive B cells.** (A) Circos plots obtained from VH gene sequences isolated from mice 14 days following infection with $10^8$ *C. rodentium.* (B) Flow cytometry analysis of IgA-positive *C. rodentium*-positive B cells isolated from Peyer patches. (C) Frequency of CD19-positive CD4-positive and CD8-positive lymphocytes in spleens of infected mice.

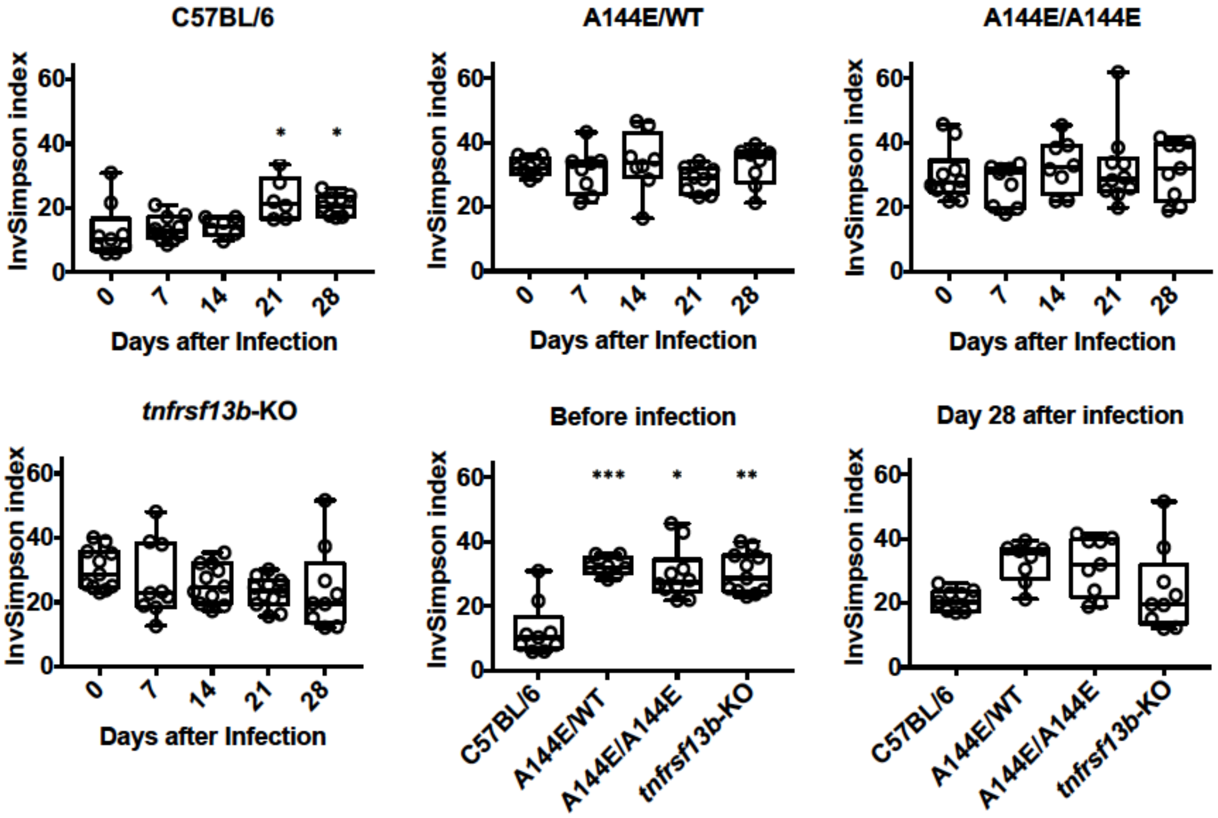


Figure S6

**Fig.S6. Microbiota diversity.** Species diversity measured using the InvSimpson Index. Measurements were determined before and weekly after infection with *C. rodentium*. Figure shows that *Tnfrsf13b*-mutant mice have greater microbiota diversity compared to C57BL/6 mice before infection but not after. Diversity only changes in C57BL/6 mice following infection. Analysis comparing diversity in *Tnfrsf13b*-mutant mice and C57BL/6 mice was by the Kruskal-Wallis test. p<0.05 *, p<0.01**, p<0.001***.

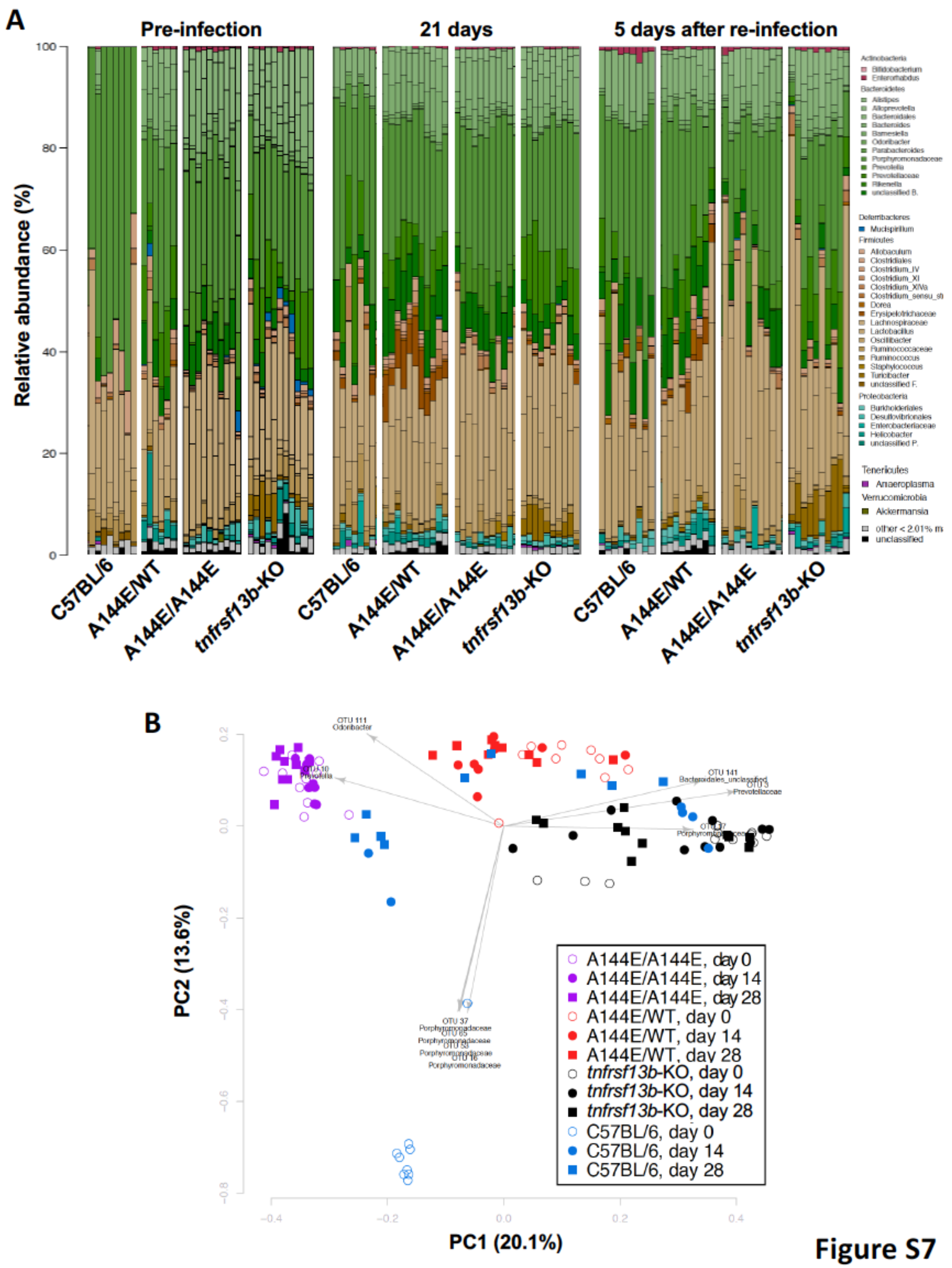


**Figure S7**

**Fig.S7. Microbiota composition.** (A) Schematic of the species composition in mice before and after *C. rodentium* infection. (B) Principal Component Analysis of microbiota composition at different times after infection. Analysis shows that microbiota in *Tnfrsf13b*-mutant mice remains clustered throughout infection while microbiota in C57BL/6 mice change dramatically.
